# Genomic predictions of climate change vulnerability in the emblematic mountain butterfly *Parnassius apollo*

**DOI:** 10.64898/2026.06.22.733620

**Authors:** Thomas Francisco, Flora Lambert-Auger, Grégory Mazoyer, Laurence Després

## Abstract

The unprecedented rate of climate warming threatens many species, and assessing their vulnerability to climate change represents a critical challenge in conservation biology. The Apollo butterfly, an emblematic mountain species, is expected to be impacted by climate change. Here, we analysed thousands of SNPs from 101 localities across Apollo French distribution. We identified 93 SNPs strongly associated with climate variation using five genotype-environment association analyses. We forecasted future climate maladaptation of French Apollo populations using four genomic offset methods and integrated these results with neutral and adaptive genetic diversity, genetic structure and adaptive climatic niches to infer their vulnerability to climate change. Jura and Alps populations exhibited the lowest risk of vulnerability to climate change, with low genomic offsets, high genetic diversity and connectivity, whereas Auvergne populations showed the highest genomic offsets and lowest neutral and adaptive genetic diversity. Only a reduced percentage (<1%) of the current distribution is predicted to face climatic conditions outside the current range, suggesting that adaptive variability required to adapt to future climates may already be present, and that assisted gene flow could represent an effective conservation strategy. Finally, we discuss some of the main challenges of genomic forecasts, particularly for declining non-model species.

## Introduction

Climate is a main selective factor shaping the biological organisation of organisms at multiple levels, from broad patterns of species richness along climatic gradients (*e.g.*, latitudinal gradients; 1), to fine-scale genetic differentiations underlying adaptive traits among populations exposed to contrasting environments (*i.e.*, local adaptation). A long history of experimental and genetic studies has revealed that local climate adaptation is widespread in natural populations. Butterfly populations, for example, frequently exhibit phenotypic differentiation for fitness-related traits along temperature clines (2–6). However, contemporary anthropogenic activities are altering the climate at an unprecedented pace (7–9). These rapid changes expose species and populations to novel environmental conditions, creating a mismatch between well-adapted phenotypes and new environmental conditions, which can result in fitness reduction (*i.e.*, maladaptation; 10,11). In response, organisms can either track their ecological niches through migration, adapt to the new environmental conditions or face negative consequences that could ultimately lead to extinction (12,13). Accumulating evidence indicates that, for many species, migration rates are often slower than the pace imposed by climate change (e.g., 14,15), and population adaptation can lag behind shifting climatic conditions (e.g., 16,17). Therefore, developing reliable methods to study and predict population maladaptation is crucial to better understand and mitigate the adverse effects of climate change on biodiversity.

Next-generation sequencing has paved the way for the development of new methods to study the effects of climate change on species, and to predict their potential climate maladaptation under changing climates (e.g., 18). To forecast the extent of maladaptation that individuals or populations may experience under changing environmental conditions, Fitzpatrick and Keller (19) introduced the concept of genomic (or genetic) offset. Assuming that individuals are adapted to their climate of origin, this approach quantifies the genetic distance between their current genomic composition and that required to maintain climate adaptation under alternative environments (e.g., future climatic conditions; 20). Several methods have been developed to compute genomic offset, including for instance redundancy analyses (RDA; 21), latent factor mixed models (LFMM; 22) and gradient forest (GF; 23). Recent work has clarified the assumptions of this approach (24,25), evaluated them through simulations (26,27) and empirical studies (28,29), and proposed theoretical frameworks (30), collectively demonstrating its strong potential to forecast climate-driven maladaptation. The genomic offset approach has been extensively applied to a wide range of study systems in the recent literature (e.g., 31-36), but remains to date, unapplied to insect species in a conservation context. Furthermore, several studies have developed or integrated tools combining genomic offset with other metrics, such as neutral and adaptive genetic diversity, landscape connectivity or genetic load (37–39), to provide a more comprehensive assessment of population vulnerability, which can be defines as ‘the degree to which a system is susceptible to, and unable to cope with, the adverse effects of climate change’ (40–42).

There is growing evidence of alarming demographic declines in insect species, which may exceed those observed in other taxonomic groups (43). For example, Hallmann and colleagues (44) documented a 75% decline in insect biomass over more than 60 sites in Germany between 1989 and 2016. In the United States, data from more than 76,000 surveys across 554 species revealed that the total abundance of butterflies has decreased by 22% over the past two decades (45). There is also emerging evidence of direct impacts of climate change on insect species in the form of population declines and shrinking ranges (46–48). In this context, genomic metrics such as genomic offset could provide valuable information to help characterise and predict this vulnerability, subsequently guiding conservation actions (43). However, a recent literature search revealed a lack of systematic genomic characterisations of insect species’ vulnerability to climate change, with for example, no applications of genomic offset in a conservation context (but see 49,50 for insect pest invasiveness). The Apollo, *Parnassius apollo* (Lepidoptera: Papilionidae), is an iconic mountain butterfly species occurring in small, fragmented populations widely distributed across Eurasia (51). In France, it occurs mainly in mountainous areas between 600 and 2,400 metres above sea level. This univoltine species overwinters at the egg stage. The larvae primarily feed on *Sedum* spp., and adults are active from May to September (52). There is evidence that *Parnassius* butterflies can adapt to their local climate, with phenotypic differences observed between populations along climatic gradients. For example, evidence of local adaptation to colder environments through wing melanism has been found in *Parnassius phoebus* (53), and genes involved in energy metabolism or cellular homeostasis were differently regulated in high versus low-altitude populations of ten *Parnassius* species in China (54), while variations in diapause-linked genes expression along climatic gradients have been documented in *Parnassius glacialis* (55). Given the extensive Eurasian distribution of *Parnassius apollo* and its presence across diverse climatic conditions within the study area (i.e., all French mountain massifs), local climate adaptation is likely to occur among populations. The Apollo butterfly has experienced a rapid demographic decline over the past 70 years, leading to assessments of its conservation status at different spatial scales: ‘Near Threatened’ in Europe by the IUCN, ‘Least Concern’ nationally in France, and Vulnerable’ to ‘Endangered’ in some French regions such as Aquitaine or Auvergne. The species is also protected under the Annex IV of the Habitats Directive and listed in Appendix II of the Convention on International Trade in Endangered Species of Wild Fauna and Flora (CITES). This decline can be attributed to a combination of factors, including habitat loss, genetic erosion and long-term climatic changes (56). Mountain environments and their associated biodiversity are particularly sensitive to rapid climatic changes (16,57). Examples of local extinctions of Apollo due to early spring-late frost events have been documented in France, such as on the Causse of Larzac in the 1980’s (58). Given the high susceptibility of both mountain and insect species to climate change, together with the recent decline of the Apollo, we hypothesised that this species may be particularly vulnerable to ongoing abrupt climate change.

The main objectives of this study were twofold: (i) to explore patterns of genetic variation along climatic gradients to identify climate-associated genetic markers and gain insight into the local climate adaptation of Apollo in France, and (ii) to assess the potential vulnerability of Apollo to future climatic conditions across French mountains. To achieve these objectives, we analysed genomic data from 321 georeferenced individuals belonging to 101 localities, covering most of the Apollo range in France. First, we examined genetic differentiation patterns across sampled localities. We then investigated the drivers of genetic differentiation and searched for climate-associated loci using genotype-environment associations (GEA) methods. Finally, we forecasted the potential future climate maladaptation across the Apollo range in France by computing genomic offsets using four different methods, defining adaptive climatic niches and integrating additional information on adaptive potential based on neutral and adaptive genetic diversity to infer their vulnerability to climate change.

## Materials and methods

### Sampling and genotyping

Sampling was carried out over the course of four flight seasons (2018, 2021-2023). A total of 321 butterflies were sampled across 101 localities distributed in the main French mountain massifs: Ardeche (2), Auvergne (6), Cevennes (3), Jura (8), Northern Alps (19), Pyrenees (18) and Southern Alps (45; Figure 1; Supplementary File). To minimise the impact of this study on Apollo populations, a single leg was collected from each butterfly for DNA extraction, preferably from males at the end of the flight season (59). All sampling was conducted under permits issued by the French authorities. Genetic data were obtained using the double-digest restriction-site associated DNA sequencing method (ddRADseq; 60) using two enzymes (*SbfI* and *MspI*). Libraries were produced as described in Kebaïli et al. (61) and paired-end sequences (2 x 125 bp) were obtained on an Illumina Novaseq. The raw reads were quality-filtered and trimmed to remove adapter sequences using *BBduk* from BBmap v38.25 software (62). Cleaned reads were demultiplexed using *process_radtags* from the Stacks v2.68 (63), specifying *SbfI* and *MspI* as restriction enzymes. Reads were trimmed to 110 bp, mapped using *bwa-mem* from bwa v0.7.17 (64) to the *P. apollo* CAJQZP01 reference genome (65), and filtered on their mapping quality (MAPQ>20). Locus reconstruction and single nucleotide polymorphism (SNP) calling were performed using the *gstacks* module from the Stacks v2.68 pipeline (63). A total of 208.6 million reads were mapped to the reference genome (94.7% mapping rate), with an average of 645,822 mapped reads and a mean depth coverage of 40.6 per sample. The resulting dataset contained 509,498 SNPs across 321 individuals.

**Figure 1.**
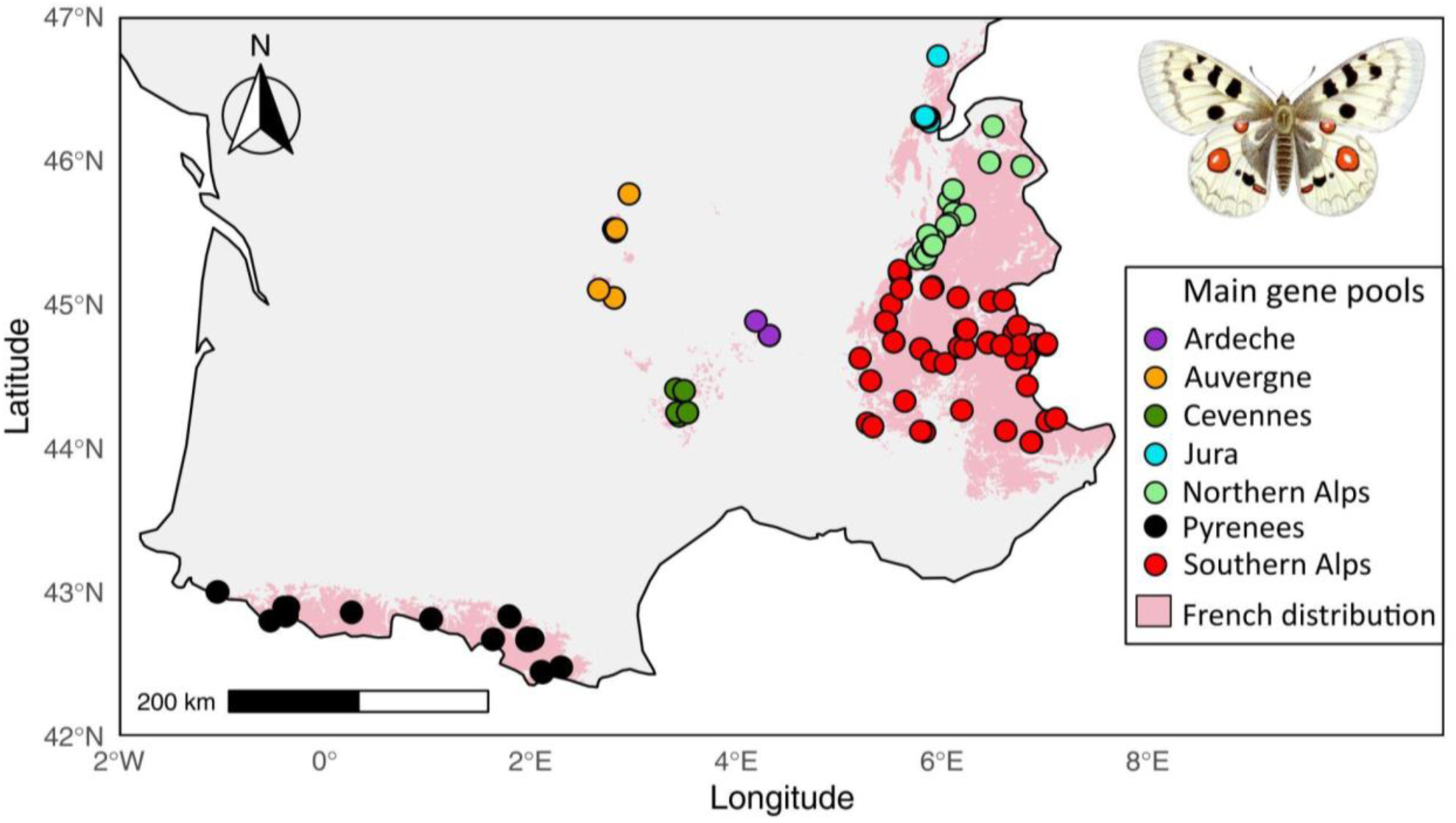
Geographical distribution of the 101 sampled *Parnassius apollo* localities coloured by main gene pools (see Figure 2), alongside its current French distribution, estimated using species distribution modelling analyses (see Methods). Illustration of *Parnassius apollo* by Jakob Hübner (95).

We then filtered out low-quality and highly physically linked SNPs (*i.e.*, 5 < depth of coverage < 150 || 0.2 < allele balance at heterozygous loci < 0.8 || linkage disequilibrium within 500bp > 0.5), producing two datasets by applying different filters to retain the most relevant variants for each analysis. First, we filtered out SNPs and individuals with more than 5% missing data, and removed SNP with a minor allele count lower than three, resulting in a final dataset comprising 17,889 SNPs across 312 individuals. This dataset (dataset 1) was used for genetic structure analyses and neutral genetic diversity estimates, since these estimates are sensitive to high rates of missing data and the removal of low-frequency SNPs (66). For GEA analyses, we filtered out SNPs and individuals with more than 30% missing data and removed SNPs with a minor allele count lower than 15 (corresponding to a minor allele frequency of 4.66%) since low-frequency alleles can lead to high false-positive rates (67). The resulting dataset (dataset 2), containing 17,131 SNPs across 318 individuals, was imputed for missing data using the most common genotype across all individuals.

### Genetic differentiation among individuals

The genetic structure among individuals was investigated by performing a principal component analysis (PCA) on the genetic data (dataset 1) using the *vegan* v2.6.4 R package (68). Additionally, the neutral genetic diversity, in the form of expected heterozygosity (*H*_S_), as well as the inbreeding coefficient (*F*_IS_) were estimated, for each distinct main gene pool, from the dataset 1 excluding the climate-associated SNPs (see below) using the R package hierfstat v0.5-11 (69; Table 1). Expected heterozygosity was also computed for the climate-associated SNPs across the main gene pools only (hereafter referred to as ‘*adaptive genetic diversity*’).

**Table 1.**
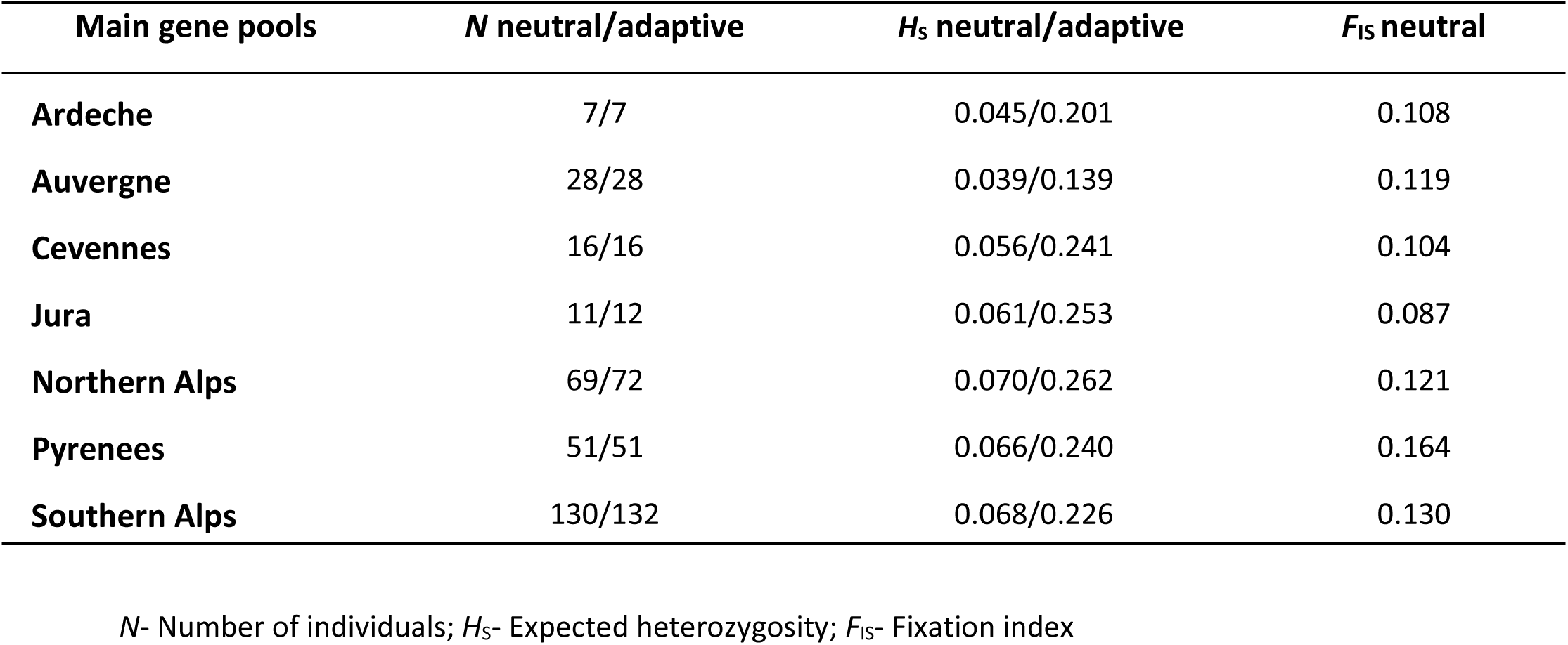
Summary statistics for the sampled Apollo individuals across the main gene pools. ‘*Neutral*’ refers to all SNPs of dataset 1 excluding the 93 climate-associated SNPs, whereas ‘*adaptive*’ refers exclusively to the climate-associated SNPs.

### Climatic data

To depict the climate under which current Apollo populations have evolved and likely adapted to, we selected the 1991-2020 period (hereafter referred to as the ‘*present*’ period). Climatic data at a spatial resolution of 30 arc-seconds (approximately 1 km² near the equator) were extracted from the Climate Downscaling Tool (ClimateDT; 70; Figure S1). This spatial resolution is adequate for Apollo since the average dispersion rate was estimated to be 400 metres using capture-mark-recapture data (51). Future conditions for the 2041-2070 period were extracted from five global climate models (GCMs; Table S1) under the AR6 scenario Shared socioeconomic pathways (SSP) 3-7.0, which reflects a strong radiative forcing and a severe climate trajectory (71). Climatic differences between current and future (mean projections of five GCMs) time periods across the 101 sampled localities (values averaged among individuals within each locality) were investigated by computing, for each main gene pool, the difference between future and current conditions (Table S3).

The identification of climatic variables potentially driving climate adaptation in Apollo populations first required selecting variables available in future climatic projections and relevant to the species’ ecology. Using this set, we performed forward selection to identify the climatic predictors best explaining the genetic variation among individuals, using the *ordiR2step* function from the *vegan* R package (68) following Francisco et al. (72) procedure. We then removed highly correlated variables (*i.e.*, Pearson’s correlation > |0.75|), retaining, when possible, the most informative predictors identified previously. Finally, we applied a threshold to the variance inflation factor to reduce the value below ten (73). The final set of predictors included: annual heat moisture index (*Annual_aridity*), mean diurnal range temperature (*Tc_mean_diurnal_range*), temperature seasonality (*Tc_seasonality*), minimum temperature of the coldest month (*Min_tc_coldest_mth*) and precipitation seasonality (*P_seasonality*; Table S2).

Present and future climatic data were extracted across the Apollo range in France, as defined by pixels from a species distribution model (SDM) with a predicted occurrence probability of at least 0.412 (see additional details in Supplementary Materials). The concordance between the climatic niche of the sampled Apollo individuals and its French distribution was confirmed using PCA based on the five climatic predictors (Figure S2).

### Evidence suggesting local climate adaptation

#### Partition of genetic variance

Following Capblancq and Forester (21), we combined redundancy analysis (RDA) and partial redundancy analysis (pRDA) to disentangle the relative contribution of geography, demographic history and climate on shaping genetic differentiation among individuals (Table 2). We ran a full RDA model with all predictors to estimate the total genetic variance explained by the model. We then ran three pRDA models to estimate the proportion of genetic variance explained by each predictor, while partialing out the effects of the others. Geography was approximated using all positive correlation axes from a spatial autocorrelation analysis (*i.e.*, distance-based Moran’s eigenvector maps) computed using the *adespatial* v0.3.23 R package (74). Historical demography was approximated using the first two axes of the genetic PCA (explaining 19.3% of the variance). Finally, the climate predictor included the five climatic variables selected previously (Table S2).

**Table 2.**
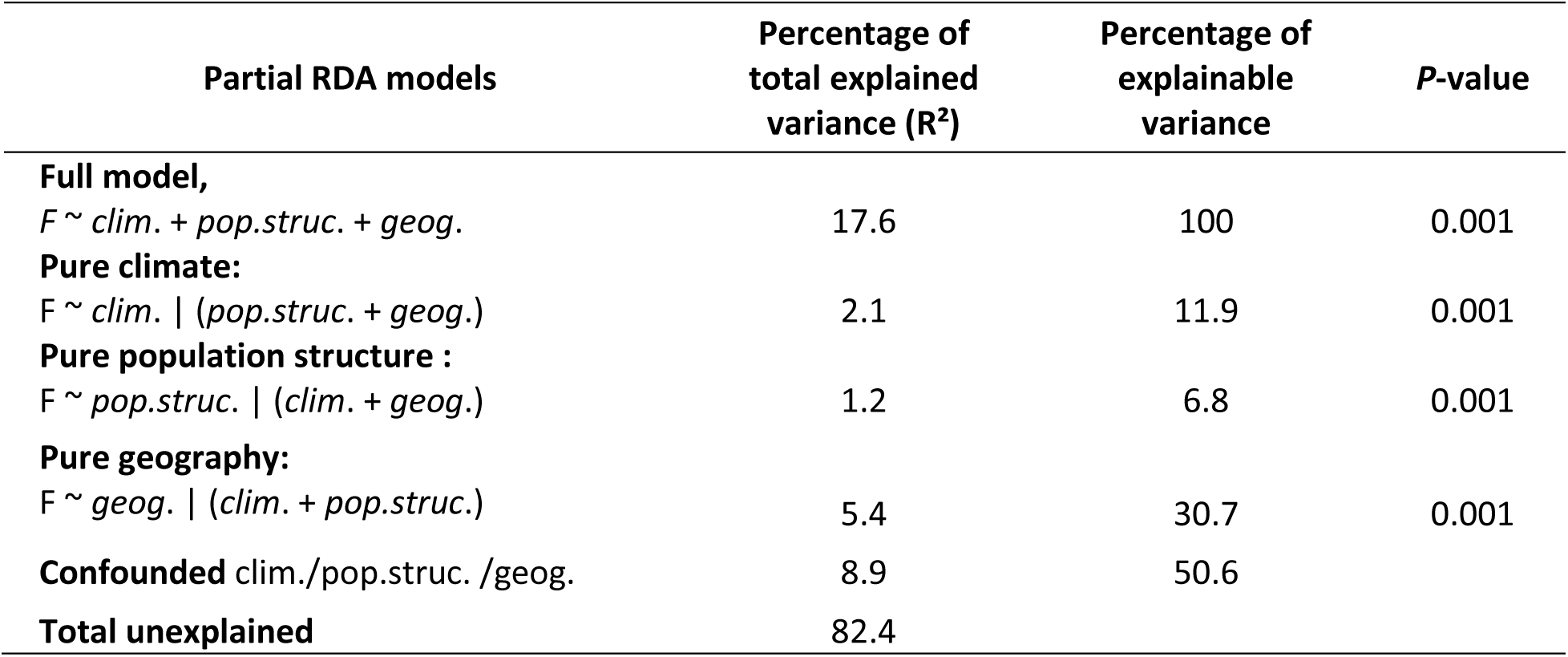
Decomposition of the total genetic variance across the sampled individuals into components attributable to climate (*clim.*), demographic history (*pop. struc.*), and geography (*geog.*), along with their confounding effects.

#### Climate-associated loci

To detect loci potentially involved, or close to genomic regions involved in climatic adaptation, five complementary GEA analyses were applied: LFMM, RDA, pRDA, and GF on uncorrected (GF_raw) and structure-corrected genetic data (GF_corrected), following Francisco et al. (72). Significant loci were selected using a Bonferroni correction of 0.01 (corresponding to a *p*-value < 5.84 * 10^-7^) or by selecting the top 1% of SNPs common across the runs. See additional information on the methods in Supplementary Materials. Finally, all SNPs identified by at least two GEA methods were defined as climate-associated SNPs.

### Potential impacts of climate change

#### Genomic offset computations

To assess the extent of potential genomic mismatch with future climate, the genomic offset of Apollo across its French range was computed using several methods (19). The general framework was similar across methods and is detailed below. Firstly, we estimated genotype-environment associations among the sampled individuals using the climate-associated SNPs and the five climatic predictors. Secondly, using these GEA models, we forecasted the genomic composition of each pixel along the French distribution for the present and future periods. Finally, we computed the genomic offset, defined for each pixel as the genetic distance between the present and future genomic compositions. Higher values of genomic offset indicate a greater genomic composition turnover needed to match the future climate, and therefore, a greater maladaptation to future climate. Genomic offset was computed using four complementary methods: two linear models that corrected for genetic structure among individuals (LFMM and pRDA; using the four latent factors defined above) and two uncorrected ones, including a linear (RDA) and a non-linear approach (GF). Additionally, for each method, we accounted for variability in future climatic predictions by interpreting the mean genomic offset values from five models run with distinct GCMs (Figure S3).

#### Extent of adaptive values

We investigated whether some of the French Apollo populations may experience climatic conditions in the future that are not currently encountered within the species’ range in France. Following Capblancq et al. (75), and using climate-associated SNPs together with the five climatic predictors, we computed two potentially ‘*adaptive enriched genetic spaces*’, one derived from RDA and the other from pRDA analyses. The latter method accounted for genetic structure by incorporating four latent factor axes derived from the LFMM analysis. For each method, we projected along the Apollo range in France the ‘*adaptive value’* of the first two canonical axes under both present and future climates. Along each axis, we then defined a ‘*present adaptive values extent*’, defined as the present range of adaptive values of Apollo across its range in France. Areas whose projected future adaptive values fell outside this present range were identified as *‘outsiders’*, meaning areas expected to face novel climatic conditions not encountered across the current Apollo range in France (34).

## Results

### Genetic and climatic differentiation among individuals

Genetic PCA revealed patterns consistent with the geographical distribution of the sampled individuals, with those from the same massif clustering together (Figure 2). We identified seven main gene pools: Auvergne, Ardeche, Cevennes, Jura, Northern Alps, Pyrenees and Southern Alps. Individuals from geographically close massifs tended to be more genetically similar. For instance, individuals from the Northern Alps and the Jura were more closely related to each other than to individuals from the Pyrenees or Auvergne. In contrast, individuals from the Cevennes and Ardeche appeared to be more genetically similar to individuals from the Jura than to Southern Alps individuals, despite the latter being closer geographically. Individuals from the Jura, Northern and Southern Alps were genetically very close, and those from the Pyrenees also clustered closely, despite extending across large geographical areas with contrasted altitudinal and climatic conditions. Neutral genetic diversity ranged from 3.9% in Auvergne up to 7% in the Northern Alps (Table 1) while adaptive genetic diversity ranged from 13.9% in Auvergne to 26.2% in Northern Alps. The five retained climatic predictors vary considerably across the seven main mountain ranges, as well as among localities within these mountains, with the highest variability being found between localities from the Southern Alps (Figure S1A). The altitude of the localities sampled in the Southern Alps ranges from 779 m to 2,410 m, while the localities in Auvergne and Jura, which show the lowest temperature and precipitation seasonalities and the lowest diurnal range, were sampled between 667 and 1,700 m. Interestingly, patterns of climatic changes between current and future periods also varied considerably across the seven main gene pools (Figure S1B and Table S3). For instance, although the Auvergne and Jura massifs currently exhibit similarly low precipitation seasonality, they are projected to experience markedly different changes, with Auvergne showing, on average, a more than twofold increase in precipitation seasonality as compared to Jura. More generally, all the climatic variables (aridity, temperature and precipitation seasonality, minimum temperature of the coldest month, and diurnal range) are projected to increase in the future for all the seven main gene pools. Some gene pools, such as the Pyrenees and Southern Alps, are expected to undergo larger changes for most predictors, while others, such as the Northern Alps, will experience smaller changes consistently across all predictors. Auvergne currently has the lowest precipitation seasonality and is expected to experience the greatest change for this climatic variable.

**Figure 2.**
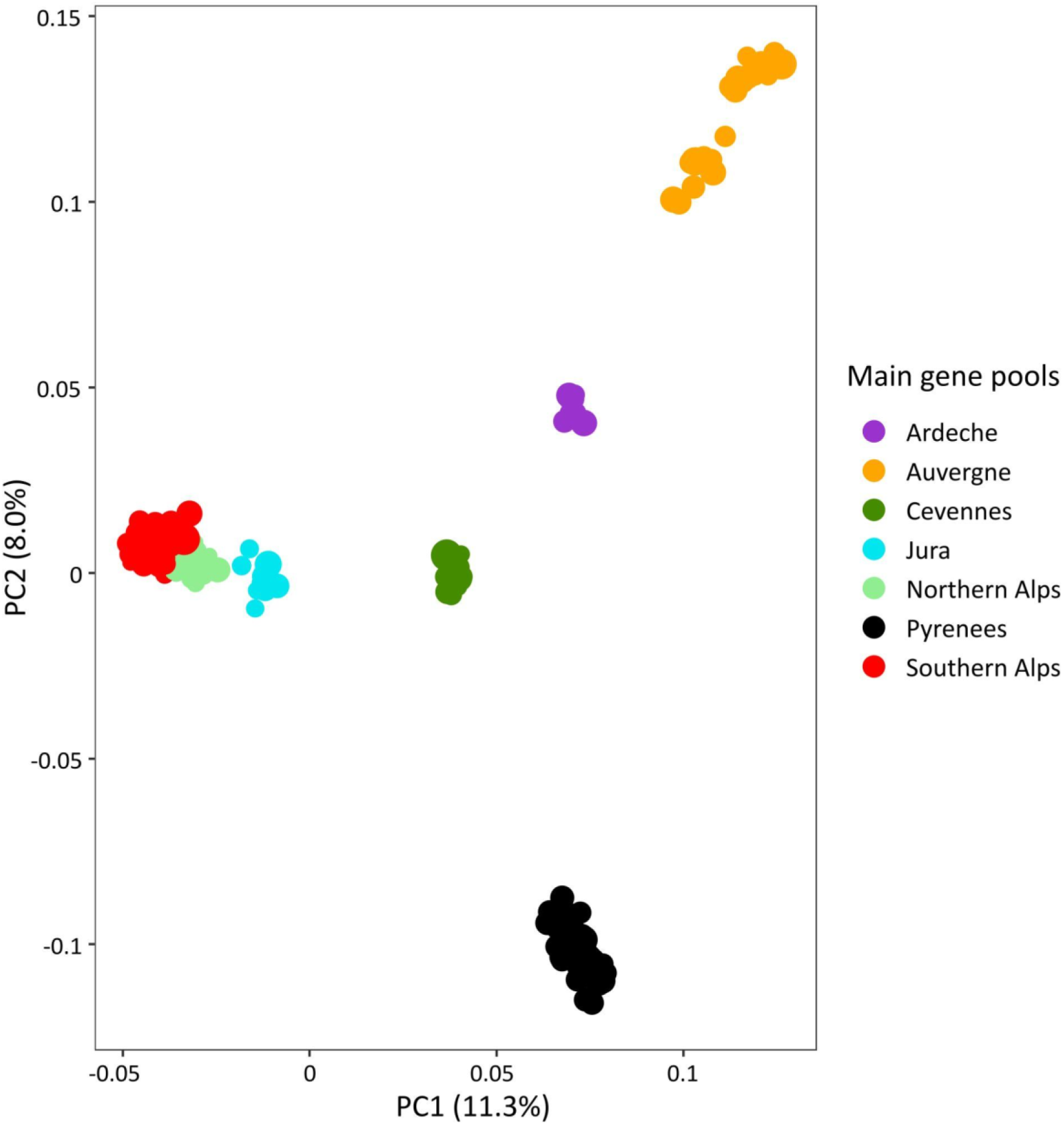
Principal component analysis (PCA) based on the genomic dataset (dataset 1), showing the position of the sampled individuals along the first two principal components. Colours refer to the main massifs of origin (hereafter referred to as ‘*main gene pools*’).

### Patterns of local climate adaptation

Variance partitioning analyses revealed that the genetic structure, geography and climate only explained a limited proportion of the genetic variance between sampled Apollo individuals (17.6%; Table 2). Among those predictors, geography explained most of the explainable genetic variance (31%), followed by climate (12%) and genetic structure (7%). Half of the explainable genetic variance remained confounded between these predictors.

RDA identified the highest number of climate-associated SNPs (253), while other methods detected fewer candidate SNPs: GF_raw (160), LFMM (132), pRDA (101), and GF_corrected (90; Figure 3). The number of overlapping candidates varied between pairs of methods, but 93 loci (around 15% of the total number of candidates; hereafter referred to as ‘*climate-associated SNPs*’) were identified by at least two GEA methods. This set of SNPs was used for subsequent analyses.

**Figure 3.**
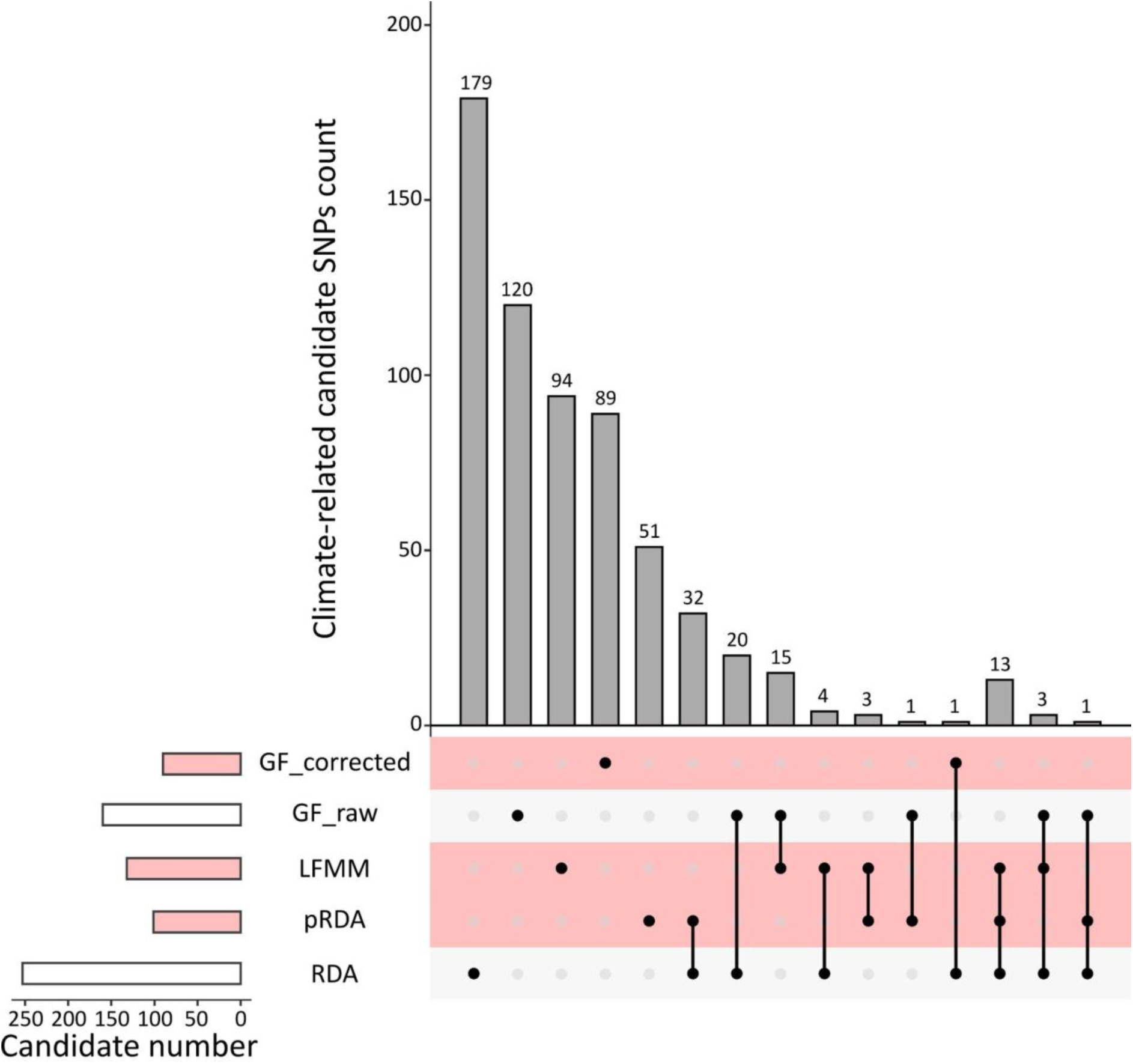
UpSet plot representing the climate-related SNPs identified by the five GEA methods. The horizontal bars display the number of candidate SNPs identified by each method, whereas the vertical bars represent SNPs unique to a given method or shared by several methods. The connected dots below indicate the contributing method(s). Colours refer to whether the method corrects for genetic structure (red) or not (white).

### Potential impacts of climate change on Apollo across its range in France

#### Genomic offset predictions

Genomic offsets computed across the sampled individuals varied across GCMs, but the extent of variation differed between methods (Figure S3). GF genomic offset predictions were consistent across GCMs, as evidenced by the high mean Pearson’s correlation coefficient (0.89). Conversely, LFMM, RDA and pRDA showed lower consistency in genomic offset predictions across GCMs (mean Pearson’s correlation coefficient: LFMM= 0.69; RDA= 0.63 and pRDA= 0.53). Importantly, genomic offset predictions were consistent across methods, as shown by the high values of all Pearson’s correlation coefficients between pairs of methods, equal or greater than 0.63 (Figure S4). Consequently, projections of the genomic offsets throughout the current range of Apollo in France were mostly consistent across methods (Figure 4). For instance, all models projected a lower genomic offset in populations from the Jura, Northern Alps and most of the Southern Alps, and a higher genomic offset in populations from Atlantic Pyrenees and Auvergne. However, some inconsistent predictions were observed across methods, such as for the populations from the central Pyrenees, which were predicted to have a moderate to high genomic offset by LFMM, RDA and pRDA methods, but a low genomic offset by GF method. Similar inconsistencies were observed for the Cevennes and Ardeche populations, for which RDA, pRDA and GF predicted a low genomic offset, while LFMM predicted an intermediate to high genomic offset.

**Figure 4.**
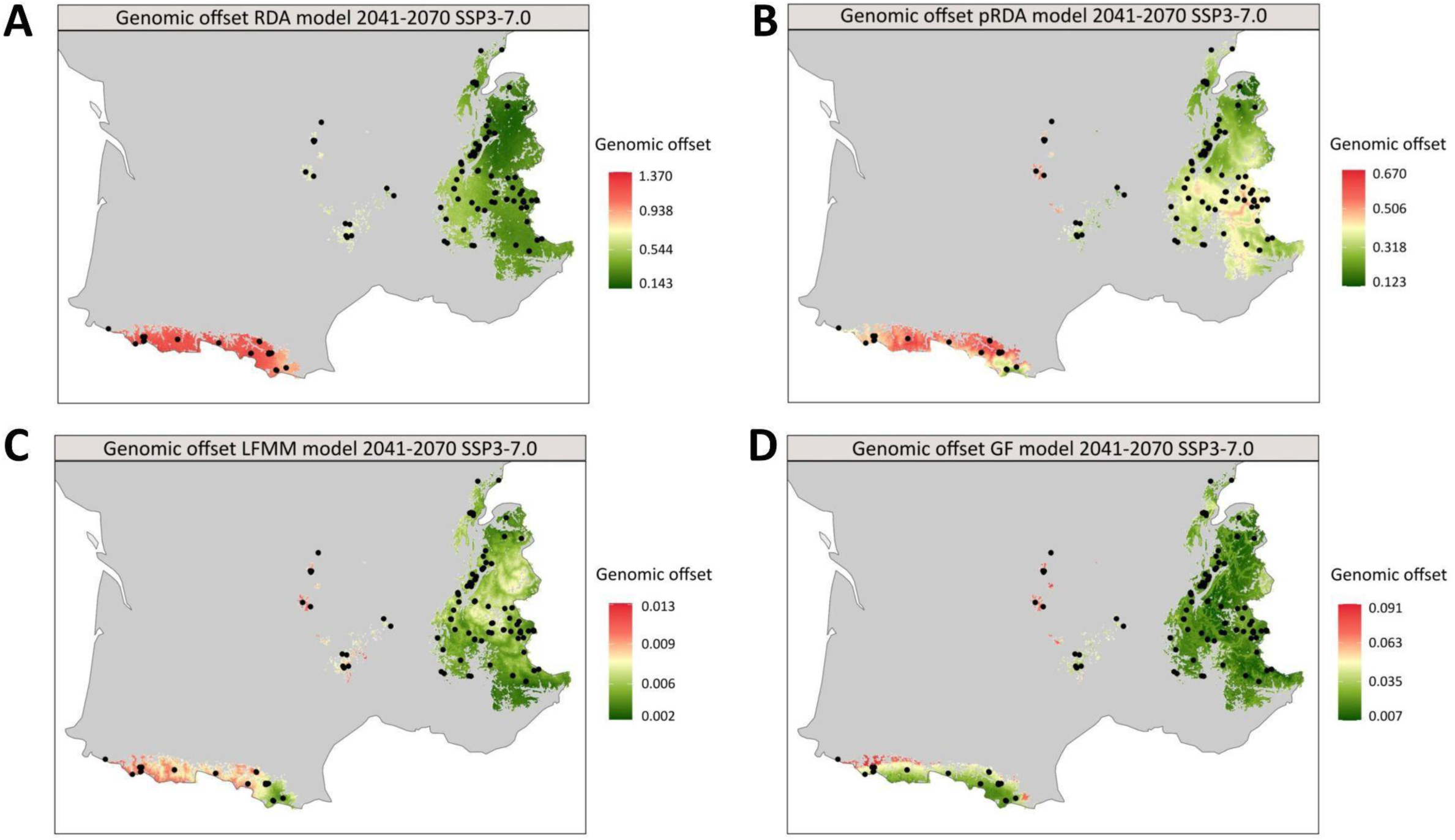
Forecasts of the genomic offsets across the Apollo range in France using four methods: RDA (**A**), pRDA (**B**), LFMM (**C**), and uncorrected GF (**D**). For each method, predictions were obtained using the 93 climate-associated SNPs and averaged from five models run with distinct general circulation models for the 2041-2070 period under the SSP3-7.0 scenario. Values range from low (green) to high (red). Black dots represent the sampled individuals used to build the models.

#### Populations facing novel climatic conditions

Precipitation seasonality and annual aridity were identified as the main drivers in the pRDA (corrected) model, whereas diurnal temperature range, temperature seasonality and minimum temperature of the coldest month were the main drivers in the RDA (uncorrected) model (Figures 5A and S5A respectively). Nevertheless, the first two retained axes of adaptive genetic spaces remained similar across the two models. Both models showed that most of the Apollo range in France is unlikely to experience future climatic conditions outside the present adaptive enriched genetic space (Figures 5B and S5B). Only a very restricted proportion of the range (0.16% for the pRDA and 0.27% for the RDA models) is predicted to encounter such novel conditions (Figures 5B, S5B and S6). Outsider regions, areas exposed to future climatic conditions not currently experienced across the Apollo range in France, are identified by both models in the Jura and the Northern and Southern Alps (Figure S6).

**Figure 5.**
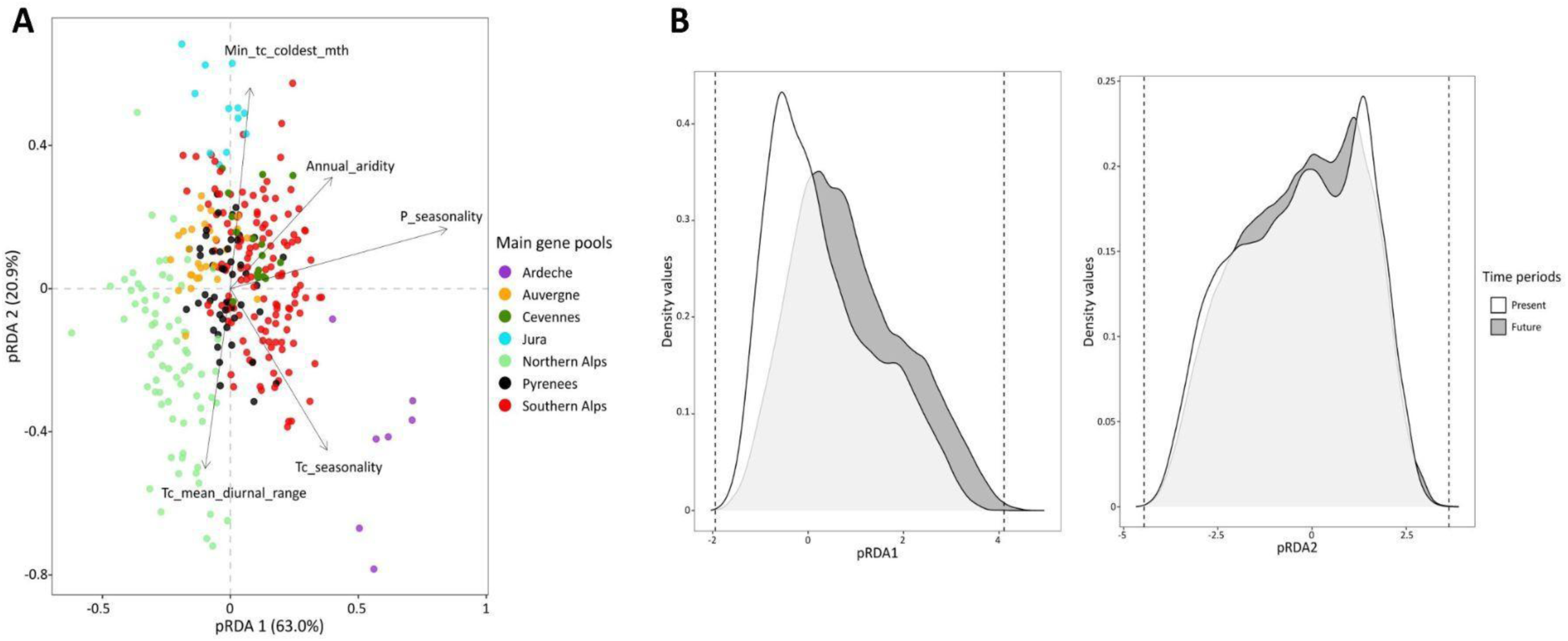
**A** Projection of the sampled individuals (points) and the five climatic predictors (arrows) onto the first two canonical axes of the pRDA adaptive climatic space (correcting for genetic structure; see Figure S5A for the uncorrected RDA). Colours refer to the main gene pools. **B** Density plots show current (white; 1991-2020 time period) and future (grey; 2041-2070 time period, based on the mean projections of five general circulation models under the SSP3-7.0 scenario) adaptive values along the first two axes of the pRDA adaptive climatic space (see also Figure S5B for the uncorrected RDA). Dotted lines delineate the present range of adaptive values (hereafter referred to as ‘present adaptive values extent’).

## Discussion

### Genetic isolation of Apollo across the French mountain massifs

The genetic differentiation of the Apollo butterfly was strongly structured by geography. Seven genetic clusters corresponding to distinct mountains were identified, and individuals from the same mountain area were found to be more genetically similar than individuals from other regions. However, individuals from the Cevennes and Ardeche were genetically closer to those from the Jura than from the Southern Alps, despite the latter being closer geographically. This suggests that the Rhône Valley represents a main barrier for this butterfly, and that the overall geographical structure observed may not solely be explained by isolation by distance. Accordingly, Kebaïli et al. (61) found that the genetic differentiation of Apollo in the Alps and Auvergne was poorly explained by isolation by distance, but rather by complex historical demography and ongoing, albeit reduced, gene flow between Alpine massifs. Several studies on mountain species have also shown that complex recolonisation scenarios of high-elevation habitats by distinct refugial lineages after glacial periods could be a key factor in explaining such genetic differentiation (76–78). Furthermore, barriers to gene flow across mountain ranges, such as large valleys and habitat fragmentation, likely contribute to strengthening the genetic differentiation observed among high altitude contemporary populations (78–81). This appears to be the case in Massif Central where relictual populations are found on the highest summits in Auvergne, Ardeche and Cevennes, that presumably diverged from each other primarily due to genetic drift. In contrast, no genetic structure was observed throughout the Pyrenees, while the Jura, Northern and Southern Alps were only weakly structured, suggesting large populations and gene flow across populations within each area, despite their large altitudinal range and wide ecological conditions. Accordingly, the highest neutral genetic diversity was found in the Alps and Pyrenees, while the lowest was found in Auvergne and Ardeche (Table 1). The fixation index (*F*_IS_) was positive in all the seven genetic clusters, indicating some substructure between samples within each mountain range.

### Evidence for an association between genetic and climate variation

Despite the main role of geography and demography in explaining the genetic variation of Apollo populations, our results support the presence of local climate adaptation in Apollo within the French mountains. The five retained climatic predictors explained a reduced, yet significant part of the observed genetic variation (2.1%, *p*-value < 0.001). Furthermore, we identified a large number of SNPs, 93, associated with the climatic predictors, despite applying conservative thresholds (i.e., Bonferroni correction of 0.01 or the top one percent of the SNPs) and approaches (i.e., retaining only SNPs identified by at least two GEA methods). The large number of candidate SNPs, along with the significant proportion of genetic variance explained by climatic predictors, suggests local climate adaptation across French Apollo populations, likely involving polygenic architectures and/ or an extensive number of traits. Accordingly, previous works have shown that climate adaptation to high-altitude in several *Parnassius* species was of polygenic nature, involving many pathways (54,55). Approximately half of the genetic variation could not be partitioned between climatic predictors and spatial genetic structure, revealing the confounding effects of geography and climate on genetic variability. Moreover, a large proportion of the genetic variance (> 80%) remained unexplained, suggesting that additional factors, such as extreme climatic events, habitat structure, or biotic pressures like pathogens or host plants, are likely to also play an important role in shaping genetic variation among French Apollo populations.

### The future of Apollo populations in France

Combining information on the spatial genetic structure, the adaptive potential, estimated from genetic diversity at neutral and climate-associated loci, and future climate maladaptation, as inferred from genomic offsets and populations falling outside the current climatic range (i.e., *outsider* populations), provided valuable insights into the vulnerability to climate change of the endangered *Parnassius apollo* butterfly across the French mountains. Most populations located in the Jura, Northern and Southern Alps share similar genetic characteristics suggesting lower risk of vulnerability to future climate. All genomic offset models forecast low to moderate climate maladaptation in these regions. In addition, substantial neutral genetic diversity at the massif level was observed (expected heterozygosity ranging from 6.1 to 7.0%), and weak spatial genetic structure, which suggests large populations with extensive gene flow, particularly between the Northern and Southern Alps. Together with the high climatic variability both within and among these regions and subsequently high adaptive genetic diversity, these results suggest that populations may already harbour the standing genetic variation, or may receive it through gene flow, to adapt to climate change. However, for some of these populations, projected to experience novel climatic conditions not currently found across the French massifs, adaptation via existing adaptive genetic variation may not be feasible.

In contrast, most genomic offset models predict that Auvergne populations will experience significant maladaptation to future climate conditions. These populations are also predicted to experience the greatest change in precipitation seasonality. Results also suggest that Auvergne populations may be small and subject to high genetic drift, as indicated by the low neutral genetic diversity and large dispersion of individual genotypes from Auvergne in the PCA as compared to larger geographical regions like the Pyrenees or the Alps. Altogether, these results emphasise that Auvergne populations are highly vulnerable to ongoing climate change, and that persistence through adaptation is unlikely to be sufficient. Upon more extensive analyses, conservation strategies such as assisted gene flow (82) could be considered for these populations, as some regions may already experience the climatic conditions Auvergne populations are projected to face in the future, thereby helping to mitigate the negative impacts of climate change. Other regions like the Pyrenees (especially in the Atlantic region) are also projected to undergo the largest temperature seasonality shifts and highest genomic offset in the future, yet they currently exhibit high genetic diversity and populations appear well connected within the region, which may help buffer these changes without the need for human interventions.

Finally, the future of several regions under climate change remains uncertain, either because different methods yield contrasting predictions of climate maladaptation (e.g., the Cevennes) or because regions exhibit low predicted climate maladaptation but suffer from reduced neutral genetic diversity and high genetic isolation (e.g., the Ardeche).

### Challenges in applying genomic offset to (endangered) non-model species

The genomic offset approach has gained considerable interest in the recent literature as a powerful tool to forecast future climate maladaptation (24–30). However, estimating genomic offset remains challenging due to several sources of uncertainty, including future climate projections and differences in model assumptions and performance. This is even more challenging for non-model species, which typically lack validation datasets. More systematic testing and reporting across diverse taxa would help improve our understanding of these limitations and enhance the robustness of predictions. Here, we observed a substantial impact of the choice of GCMs on several methods, with only moderate correlations among predictions, as also reported in previous studies (19,29,83). These results highlight that variability in future climate projections (84) should be carefully accounted for in predictive studies, even when considering a single SSP scenario. Additionally, variability in genomic offset predictions across methods was observed, although predictions remained consistent, with a minimum Pearson’s correlation coefficient of 0.63 across methods. This level of concordance is similar to or greater than that reported in previous studies using multiple approaches (29,72). These results overall support the robustness of our climate maladaptation predictions. Nevertheless, it is important noting that no evaluation of the genomic offset models against experimental or phenotypic data, or using simulation has been performed on *Parnassius apollo* and therefore care must be taken when interpreting such predictions because the link between genomic offset and fitness has not been established.

Another challenge in genomic offset forecasting lies in the number of data available to build predictive models. This is particularly relevant for species with restricted distribution, as well as declining and endangered ones, which could paradoxically be the main target of such forecasting methods (e.g., *Araucaria araucana*, 85; *Tetracentron sinense*, 86; *Tetraena mongolic*, 87; *Parnassius apollo*, this study). Here, to address these challenges, we conducted our analyses at the individual level in order to maximise the spatial coverage of French massifs and subsequently the environmental heterogeneity, following recent recommendations (88–90). Indeed, Aguirre-Liguori et al. (89) showed using empirical data that the number of sampled sites had a greater impact on genomic offset estimates than the number of individuals sampled per population. Furthermore, Lind and Lotterhos (90) demonstrated the practical use of individual-level instead of population-level data to compute genomic offset. Using simulation data, they showed that genomic offset models built using genotypes (i.e., individual-level data) rather than allele frequencies (i.e., population-level data) provided similar or even improved model performance.

Overall, genomic offset approaches offer a promising framework to investigate potential maladaptation of biodiversity to future climate change. Current challenges, particularly for declining or range-restricted species, such as limited sample sizes, may be partially addressed through individual-based modelling approaches (90). However, important uncertainties remain, particularly regarding differences among modelling methods and their predictive reliability (e.g., 29). Empirical evaluation of genomic offset predictions using phenotypic or fitness-related data could help clarify the relationship between predicted genomic mismatch and biological responses (91). In practice, however, such validation may be difficult or infeasible for many non-model or endangered species, and may delay the use of genomic predictions in conservation planning. This highlights the need for broader assessments of genomic offset models to better define the contexts in which their predictions are most informative. In situations where empirical validation is not feasible, complementary approaches such as time series (24) or comparisons among multiple genomic offset methods may help to characterise uncertainty, with particular attention to regions where predictions converge across methods, as illustrated in Francisco et al. (72) and the present study.

## Conclusions

Genomic analyses of 321 Apollo butterflies across their distribution range in France provided valuable insights into their adaptation to local climates and vulnerability to climate change. A significant portion of genetic variability across Apollo was explained by climatic predictors, with 93 SNPs being strongly associated with these predictors. Furthermore, multiple genomic offset analyses provided consistent predictions across most of the species’ range in France. Combining these predictions with neutral and adaptive genetic diversity, genetic structure, and adaptive climatic niches information, Auvergne populations were identified as the most vulnerable to climate change, while Jura and Alpine populations appeared to be the least at risk. Notably, less than 1% of the current French Apollo distribution is expected to face novel climatic conditions in the future, suggesting that adaptive variability required to face future climate may already be present, and that assisted gene flow could be a viable conservation strategy. However, substantial unexplained genetic variance between Apollo and the presence of isolated populations in some regions highlights the need for further investigation to assess the risks of outbreeding depression and disruption of non-climatic local adaptation prior to the implementation of assisted gene flow. Finally, additional factors, whether linked to climate change or not, such as upward shifts in the tree line (92,93) and the dynamics of its host plants (see, for example, the case of the *Euphydryas aurinia* butterfly; 94) are also expected to strongly shape the future distribution of Apollo.

## Data Archiving Statement and Code Availability

The raw sequences (FASTQ files) analysed in the present study are available in the European nucleotide archive repository (ENA) at EMBL-EBI (http://www.ebi.ac.uk/ena) and accessible under study accession numbers PRJEB48121 and PRJEB110627. The VCF file obtained after pre-treatment, containing 509,498 SNPs across 321 individuals, as well as all the relevant code used in this study, are openly available in Zenodo at: https://doi.org/10.5281/zenodo.19222056.

## Author Contributions

Designed research: LD, TC, TF; obtained funding: LD; genetic data acquisition and pretreatment: LD, TF, FL-A; data analysis: TF, FL-A, GM; wrote the paper: TF, LD, FL-A. All authors critically read, edited, commented and approved the submitted version of the paper.

## Acknowledgments

We thank Gaelle Sobczyk-Moran (OPIE) and Jean-Marc Salles (DREAL Auvergne Rhône-Alpes) for administrative and financial support to this partenarial academic-wildlife management project, conducted as part of the national action plan for butterflies, including collection permits. We also warmly thank all the collectors and their respective structures: Paul Boudin, Jérôme Bailly (PNR Chartreuse), Yann Baillet, Philippe Bordet (FLAVIA APE), Hervé Tournier, Brice Palhec (RNN Hauts-plateaux du Vercors), Thierry Leroy (RNN Chastreix-Sancy), Mathilde Pantalacci, Philippe Francoz, Jean-François Lopez (PNR Bauges), Pierre Durlet (PNR Haut Jura), Frédéric Mora, Perrine Jacquot (CBN Franche-Comté), Cécile Lemarchand (RNR Gorges de Daluis), Marianne Elias, Violaine Ossola and Roland Teissier (MNHN), Mathieu Joron, Paul Doniol-Valcroze (CEFE), Jocelyn Fonderflick (PN Cévennes), Pierre Desriaux, Jean-Marie André, Stéphane Bence, Sonia Richaud (CENA PACA), Jean Philippe Reygade (SM PNR Volcans d’Auvergne), Nicolas Maurel (Proserpine), Mathieu de Lamarre (LECA), Bernard Bal (CEN Haute-Savoie), Rémi Lafitte (RN Mantet), Maxime Juignet (RN Les Partias), Damien Cocâtre (RN Ardèche), Philippe Bachelard (SHN Alcide d’Orbigny), Mathieu Molières (Cistude Nature), Philippe Loudin (RNN Vallée de Chaudefour), Lily Dunyach (FRN Catalane), David Soulet, Loyann Boy (RN Aulon, CEN Occitanie), Christophe Lhez (RN Orlu), Alexis Calard, Laurent Servières, Florine Hadjaj (ANA-CEN Ariège), and Eric Drouet (Oreina). We gratefully acknowledge the global biodiversity information facility (GBIF; accessions: https://doi.org/10.15468/dl.sks7dz, https://doi.org/10.15468/dl.35yftf and https://doi.org/10.15468/dl.d2a6su), iNaturalist, Écrins National Park, the Flavia Association, and Artemisiae for providing occurrence records of *Parnassius apollo* and *Sedum album*. This work was supported by the grant n° 2103826137 from the French government and by the computing facilities from the University Grenoble Alpes (GRICAD).

