## Supplementary Material for "Genomic predictions of climate change vulnerability in the emblematic mountain butterfly *Parnassius apollo*"

### Supplementary Materials

### *Supplementary Methods*

#### Species distribution model

The species distribution model (SDM) was built and projected using an ensemble modelling framework (1) implemented in the R *biomod2* package (2). Environmental predictors included climatic, topographic, and land-cover variables. Climatic variables were downloaded from CHELSA v2.1 (1981-2010) at a spatial resolution of 30 arc-seconds (approximately 1 km² near the equator; 3). All the variables were cropped to the study area (-2.786, 8.992, 42.268, 47.725). To reduce multicollinearity, climatic variables were selected based on Pearson's correlation coefficients (r <0.7) and the prior knowledge of *P. apollo* ecology. The following variables were selected: mean diurnal range (BIO2), isothermality (BIO3), temperature seasonality (BIO4), minimum temperature of coldest month (BIO6), mean temperature of wettest quarter (BIO8), precipitation of driest month (BIO14), precipitation seasonality (BIO15), and the annual amplitude of cloud cover (CLT range). Land cover variables were derived from the CORINE land cover database, and were binarised and grouped into six classes: natural grasslands, shrub heathlands, sparse vegetation, rocky habitats, anthropogenic areas and forests. Elevation data and solar radiation were downloaded from WorldClim v.2 (4). Elevation data was used to compute topographic variables, including terrain roughness and aspect, and solar radiation was included as an additional predictor in the SDM models. To incorporate the main Apollo larval host plant into the Apollo SDM, we retrieved 2,021 occurrence records of *Sedum album* from the GBIF database (GBIF: https://doi.org/10.15468/dl.35yftf & https://doi.org/10.15468/dl.d2a6su), and developed a *Sedum album* SDM using the same method applied for Apollo SDM (see below). The probability of host plant's presence predicted by this model was subsequently incorporated as a predictor into the Apollo SDM. To reduce spatial autocorrelation, occurrence records were spatially filtered to retain one single point per 1 km² pixel using the *spThin* v.1.3.16 R package (5).

For the Apollo SDM, we used 2,620 occurrences extracted from various databases (GBIF: https://doi.org/10.15468/dl.sks7dz; iNaturalist; Écrins National Park; Flavia association and Artemisiae) for the period 2010-2025. We built an ensemble model based on eight algorithms: generalised linear model (GLM), generalised additive model (GAM), generalised boosted model (GBM), flexible discriminant analysis (FDA), random forest (RF), random forest for unbalanced data (RFd), maximum entropy (MAXNET) and extreme gradient boosting (XGBOOST). As reliable absence data were unavailable, 10 replicates of 7860 random pseudo-absences were generated for each model. Indeed, a pseudo-absence to presence ratio of 3:1 was applied following recommendations and practical guidelines provided in the biomod2 documentation. The models were calibrated by cross-validation with five repetitions, using 70% of the data for calibration and 30% for validation. Model performance was evaluated using the Boyce index (6), which is specifically suited for assessing predictive performance of presence-only models. Only models with a Boyce index greater than 0.7 during the calibration phase were retained for the ensemble modelling. The final ensemble model was constructed using weighted mean (EMwmean), which gives higher influence to models with better performance.

A threshold value (0.412) was obtained from model predictions by identifying the probability cut-off that maximized the Boyce index during model evaluation, and was used to classify habitat suitability. Model evaluation metrics, including the Boyce index, were extracted using the '*get_evaluations*' function of the Biomod2 package.

**Climate-associated loci**

To detect loci potentially involved, or close to genomic regions involved in climatic adaptation, five complementary GEA analyses were applied: LFMM, RDA, pRDA, and GF on uncorrected (GF_raw) and structure-corrected genetic data (GF_corrected), following Francisco et al. (7).

Latent factor mixed models (LFMM; 8) use linear regression to estimate the effect of environmental predictors on each SNP, accounting for genetic structure using latent factors (four latent factors were included). Depending on the method, significant loci were selected using a Bonferroni correction of 0.01 (corresponding to a $p$-value < $5.84 * 10^{-7}$).

Redundancy analysis (RDA) and partial redundancy analysis (pRDA) are linear, multivariate approaches combining ordination and multiple regression analyses (9). In pRDA, genetic structure is accounted for using the four latent factor axes from the LFMM analysis. Candidate loci were defined, according to Capblancq and Forester (10), as SNPs whose Mahalanobis distances along the first two canonical axes were unusually high compared to the centroid of the redundancy space, applying a Bonferroni correction of 0.01.

Furthermore, we employed the gradient forest (GF) algorithm to identify non-linear relationships between allele frequencies and environmental variables (11,12). This ensemble tree-based method models allele frequency turnover along environmental gradients. We ran two types of models: one based on unadjusted genomic data (GF_raw), and another using structure-corrected data (GF_corrected), calculated by multiplying the LFMM-adjusted effect size matrix by the transposed climatic variable matrix (see 7). Each model was replicated five times, with the top 1% of SNPs common across the runs being selected as candidates.

### Supplementary Tables & Figures

*Table S1*. Description of the five general circulation models (GCMs) used to forecast genomic offset and adaptive climatic niches under the SSP3-7.0 scenario for the 2041-2070 time period.

| **Global climate models (GCMs)** | **References**[1] |
|---|---|
| GFDL-ESM4.1 | Dunne et al. (13) |
| IPSL-CM6A-LR | Boucher et al. (14) |
| MPI-ESM1-2-HR | Gutjahr et al. (15) |
| MRI-ESM2-0 | Yukimoto et al. (16) |
| UKESM1-0-LL | Sellar et al. (17) |

[1]References

*Table S2*. Description of the five climatic predictors used in genotype-environmental association (GEA) and genomic offset analyses.

| Variable | Abbreviation |
|---|---|
| **AHM: Annual heat moisture index (°C /mm)** | Annual_aridity |
| **Bio2: Mean diurnal range (°C)** | Tc_mean_diurnal_range |
| **Bio4: Temperature seasonality (standard deviation °C x100)** | Tc_seasonality |
| **Bio6: Minimum temperature coldest month (°C)** | Min_tc_coldest_mth |
| **Bio15: Precipitation seasonality (coefficient of variation)** | P_seasonality |

*Table S3*. Overview of the climate differences between the current (1991-2020) and future (2041-2070, based on the mean projections of five general circulation models under the SSP3-7.0 scenario) periods across the 101 sampled localities (values averaged between individuals within each locality). Positive values indicate an increase from the present to the future conditions.

| **Main gene pools** | **Annual aridity** *(°C/mm)* | **Tc mean diurnal range** *(°C)* | **Tc seasonality** *(standard deviation °C * 100)* | **Min tc coldest mth** *(°C)* | **P seasonality** *(coefficient)* |
|---|---|---|---|---|---|
| **Ardeche** | 0.82 | 0.33 | 48.56 | 1.97 | 0.16 |
| **Auvergne** | 1.38 | 0.44 | 51.96 | 1.45 | 2.66 |
| **Cevennes** | 1.09 | 0.42 | 52.43 | 1.34 | 1.86 |
| **Jura** | 0.90 | 0.35 | 29.15 | 1.60 | 1.15 |
| **Northern Alps** | 0.85 | 0.21 | 35.07 | 1.59 | 0.24 |
| **Pyrenees** | 2.20 | 0.75 | 68.13 | 0.73 | 0.87 |
| **Southern Alps** | 2.35 | 0.11 | 35.32 | 2.34 | 0.60 |

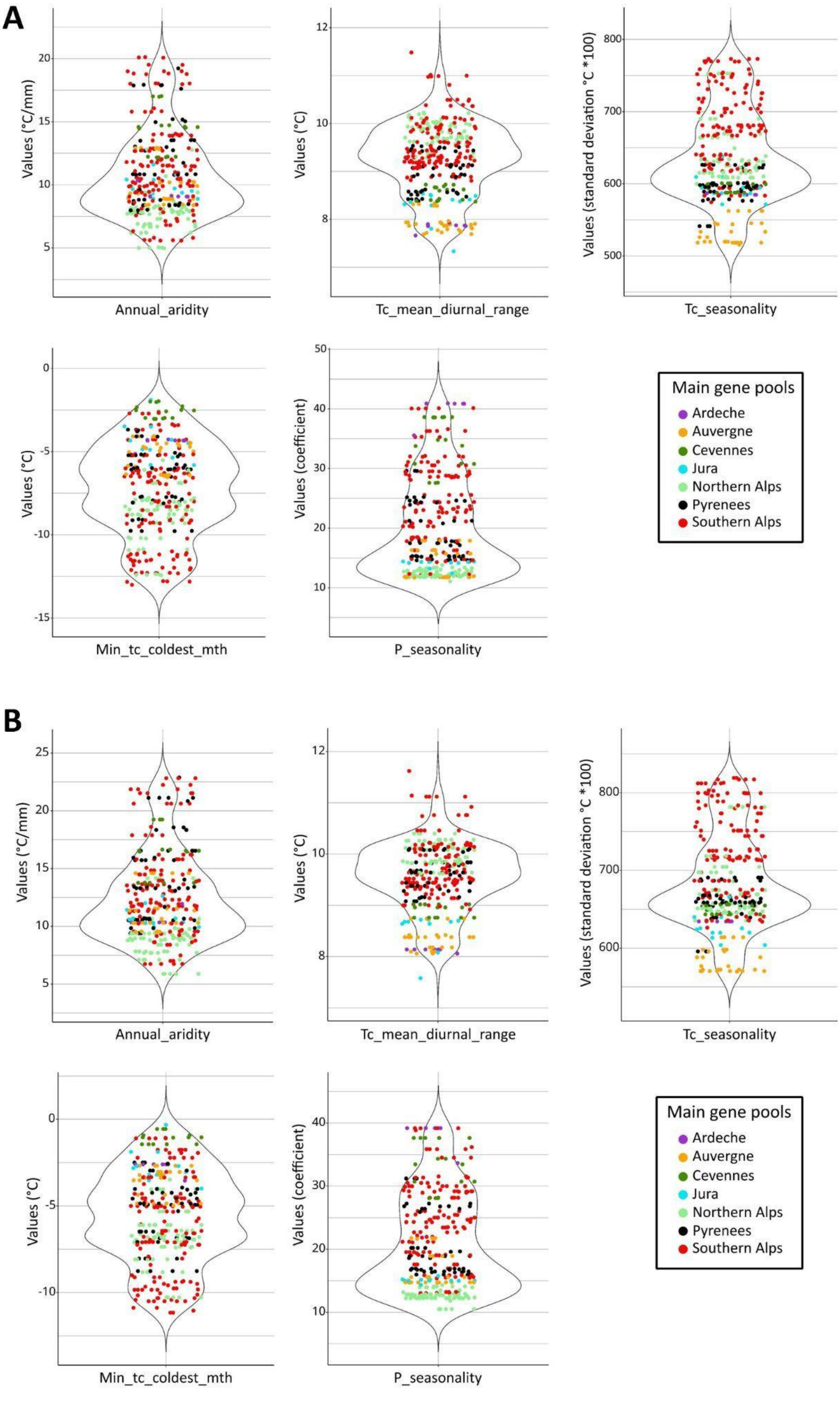
A
Values (°C/mm)
Annual_aridity
Values (°C)
Tc_mean_diurnal_range
Values (standard deviation °C *100)
Tc_seasonality
Min_tc_coldest_mth
Values (coefficient)
P_seasonality
Main gene pools
Ardeche
Auvergne
Cevennes
Jura
Northern Alps
Pyrenees
Southern Alps
B
Values (°C/mm)
Annual_aridity
Values (°C)
Tc_mean_diurnal_range
Values (standard deviation °C *100)
Tc_seasonality
Min_tc_coldest_mth
Values (coefficient)
P_seasonality
Main gene pools
Ardeche
Auvergne
Cevennes
Jura
Northern Alps
Pyrenees
Southern Alps

*Figure S1*. Violin plots displaying the climate conditions for the five retained climatic predictors for the (A) current (1991-2020) and (B) future (average values across the five general circulation models for the period 2041-2070 under the SSP 3-7.0) time periods across the sampled individuals, colours per main gene pools.

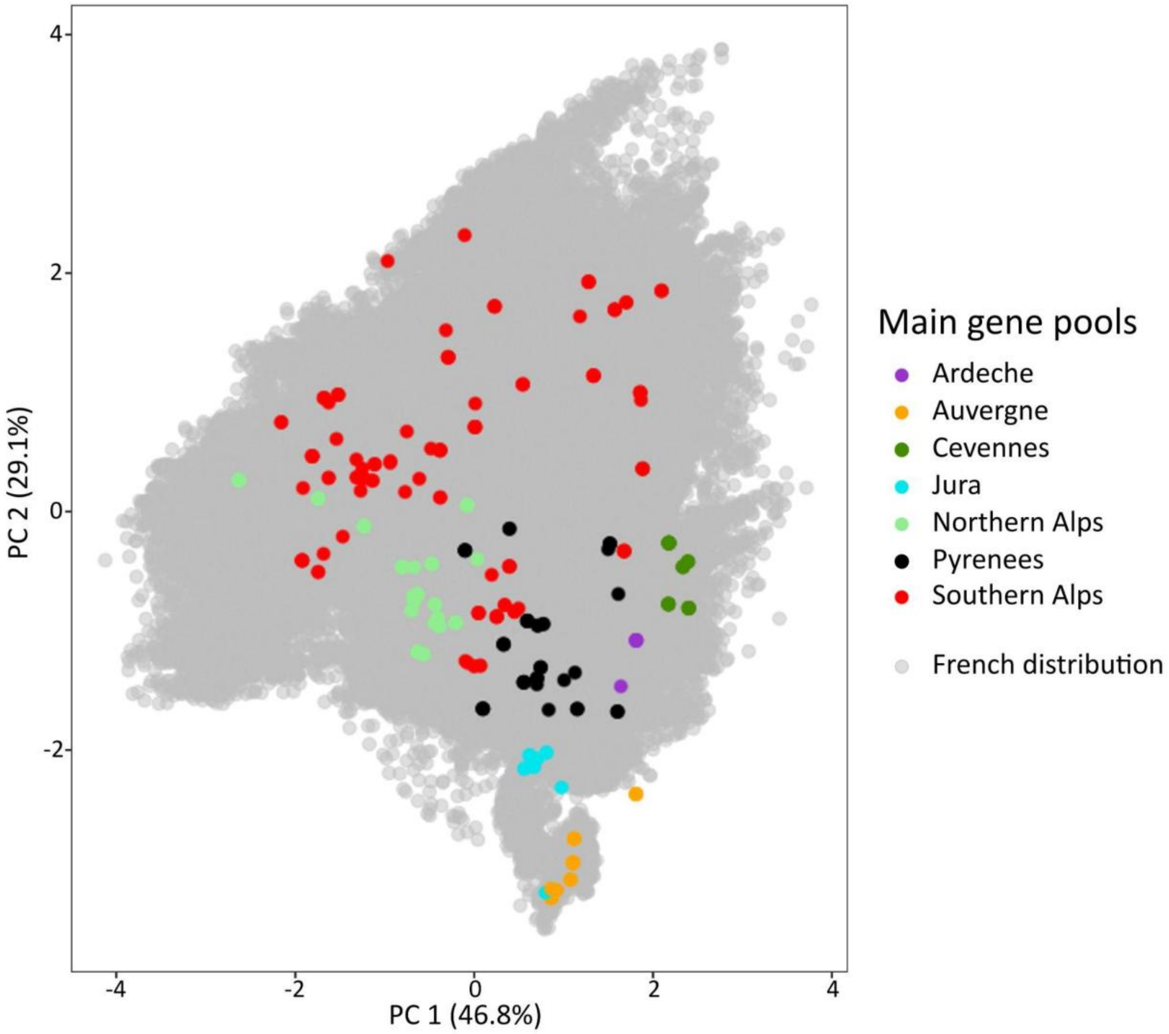


*Figure S2*. Principal component analysis showing the climatic envelope of the sampled individuals (coloured per main gene pools) against the climatic envelope of the Apollo range in France (grey; based on species distribution modelling, see Main text) for the five considered climatic predictors under the 1991-2020 time period.

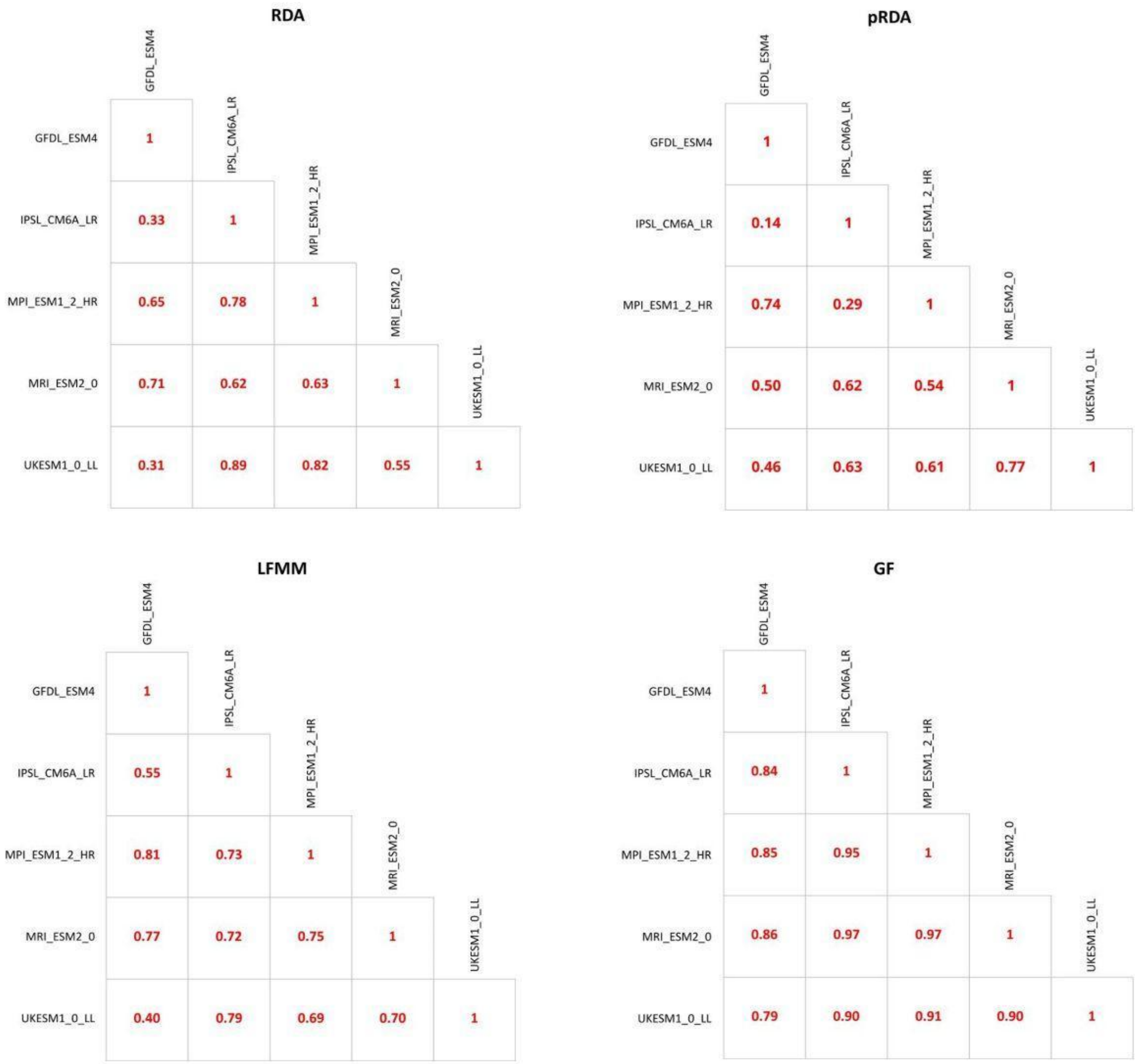


*Figure S3*. Pearson's correlation matrix showing, for each method, the correlation coefficients of genomic offset values for the sampled individuals across the five models run with distinct GCMs for the 2041-2070 period under the SSP3-7.0 scenario. All correlation coefficients are significantly positive.

| | GF | LFMM | RDA | pRDA |
|---|---|---|---|---|
| GF | 1 | | | |
| LFMM | 0.63 | 1 | | |
| RDA | 0.63 | 0.99 | 1 | |
| pRDA | 0.67 | 0.96 | 0.94 | 1 |

*Figure S4*. Pearson's correlation matrix showing the correlation coefficients of genomic offset values (forecast for the 2041-2070 time period, based on the mean projections from five general circulation models under the SSP3-7.0 scenario) for the sampled individuals across the four methods. All correlation coefficients are significantly positive.

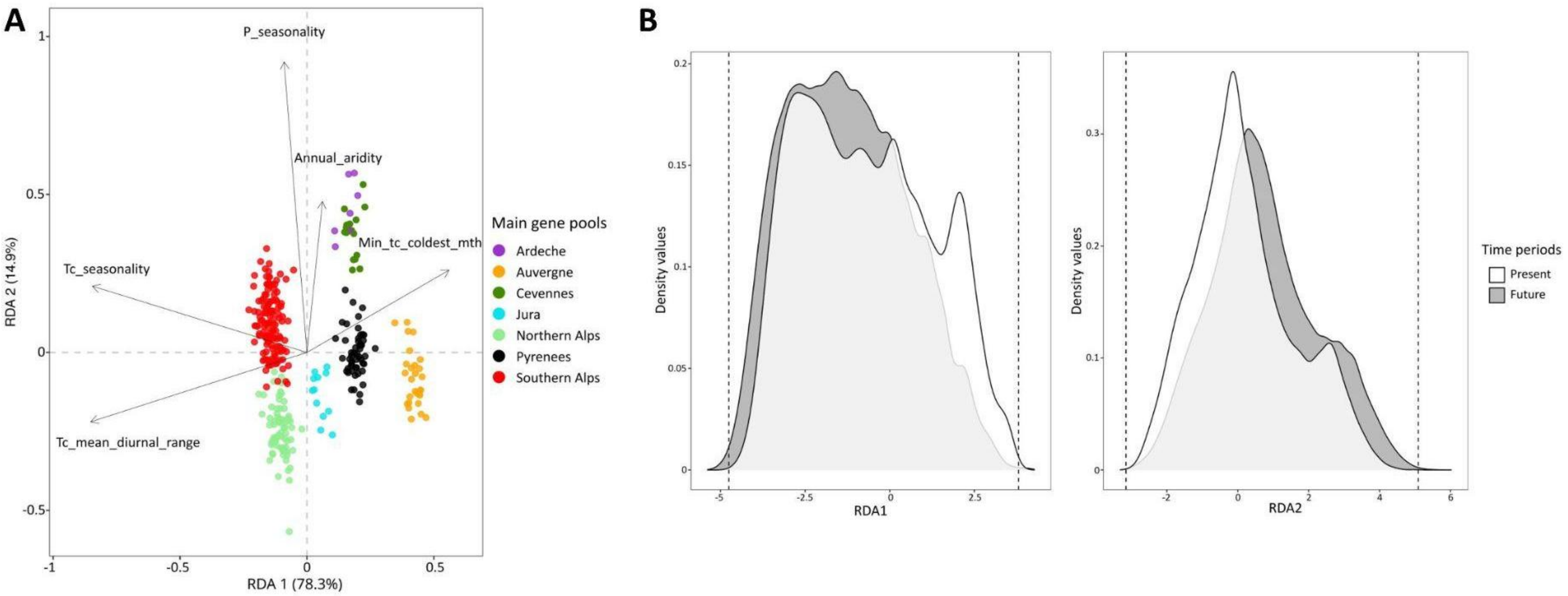


*Figure S5*. **A** Projection of the sampled individuals (points) and the five climatic predictors (arrows) onto the first two canonical axes of the RDA adaptive climatic space model (uncorrected for genetic structure; see Figure 5A for the corrected pRDA model). Point colours refer to the main gene pools. Note that the axes are swapped relative to the pRDA model presented in the main text (Figure 5), such that RDA1 approximates sign-reversed pRDA2 and RDA2 approximates pRDA1. **B** Density plots show current (white; 1991-2020 time period) and future (grey; 2041-2070 time period, based on the mean projections of five general circulation models under the SSP3-7.0 scenario) adaptive values along the first two axes of the RDA adaptive climatic space. Dotted lines delineate the present range of adaptive values (hereafter referred to as ‘present RDA adaptive value extent’).

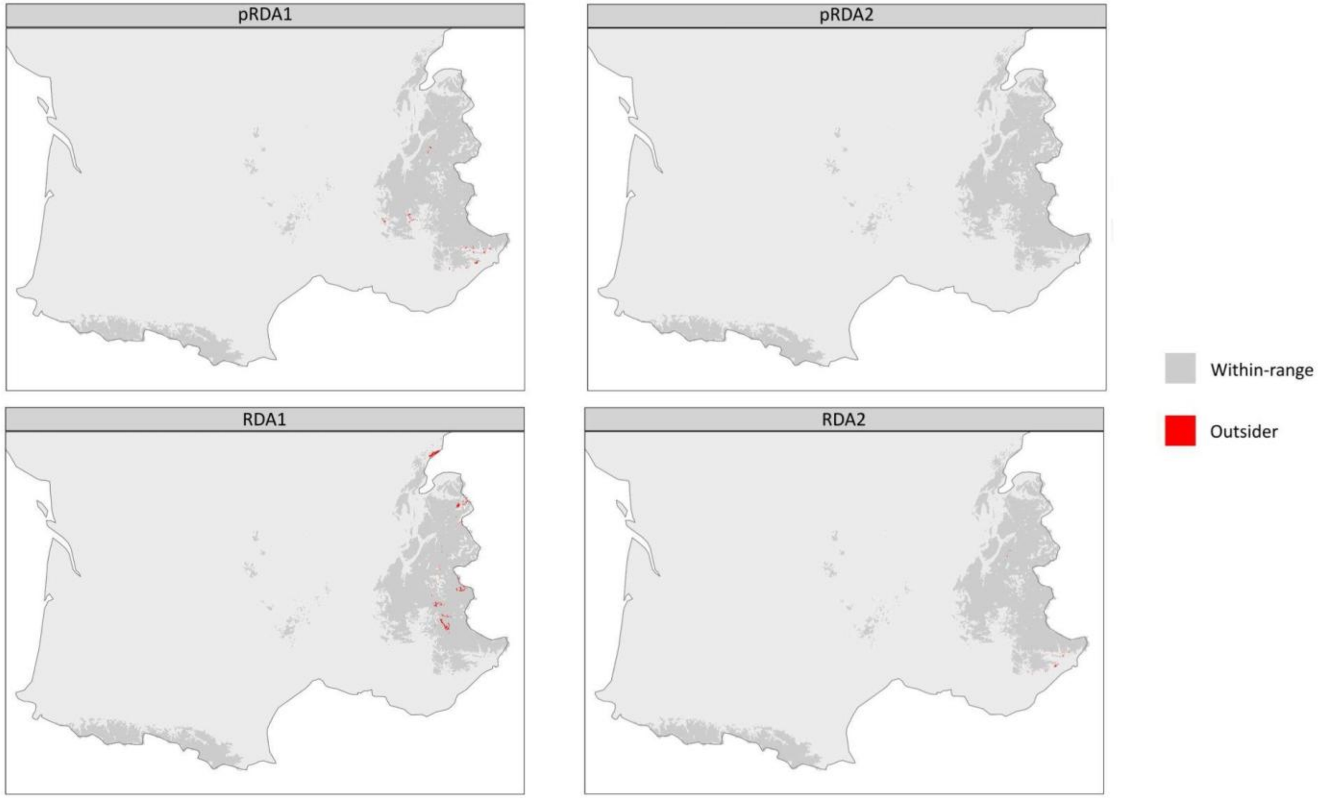


*<u>Figure S6</u>*. Projection of the adaptive values of the first two canonical axes of the pRDA and RDA adaptive climatic spaces across the Apollo range in France. Values were defined as ‘within-range’ when falling inside the present adaptive climatic space (Figures 5 and S5) and as ‘outsider’ when falling outside.
